# Functional Characterisation of the ATOH1 Molecular Subtype Indicates a Pro-Metastatic Role in Small Cell Lung Cancer

**DOI:** 10.1101/2024.02.16.580247

**Authors:** Alessia Catozzi, Maria Peiris-Pagès, Sam Humphrey, Mitchell Revill, Derrick Morgan, Jordan Roebuck, Yitao Chen, Bethan Davies-Williams, Alice Lallo, Melanie Galvin, Simon P Pearce, Alastair Kerr, Lynsey Priest, Victoria Foy, Mathew Carter, Rebecca Caeser, Joseph Chan, Charles M. Rudin, Fiona Blackhall, Kristopher K Frese, Caroline Dive, Kathryn L Simpson

## Abstract

Molecular subtypes of Small Cell Lung Cancer (SCLC) have been described based on differential expression of transcription factors (TFs) *ASCL1, NEUROD1*, *POU2F3* and immune-related genes. We previously reported an additional subtype based on expression of the neurogenic TF *ATOH1* within our SCLC Circulating tumour cell- Derived eXplant (CDX) model biobank. Here we show that ATOH1 protein was detected in 7/81 preclinical models and 16/102 clinical samples of SCLC. In CDX models, ATOH1 directly regulated neurogenesis and differentiation programs consistent with roles in normal tissues. In *ex vivo* cultures of ATOH1-positive CDX, ATOH1 was required for cell survival. *In vivo*, ATOH1 depletion slowed tumour growth and suppressed liver metastasis. Our data validate ATOH1 as a *bona fide* oncogenic driver of SCLC with tumour cell survival and pro-metastatic functions. Further investigation to explore ATOH1 driven vulnerabilities for targeted treatment with predictive biomarkers is warranted.

## INTRODUCTION

SCLC is an aggressive neuroendocrine (NE) tumour constituting ∼15% of lung cancers. SCLC is the sixth most common cause of cancer-related deaths, accounting for ∼250,000 diagnoses worldwide each year^1–4^. Most patients with SCLC present with extensive stage (ES) disease characterised by widespread metastases and rapidly acquired resistance to initially effective standard-of-care (SoC) platinum- based chemotherapy^5^. SoC was unchanged for >30 years^6^ until the recent addition of immunotherapy that extends overall survival of a minority of patients, including rare patients with durable responses^7–10^.

SCLC molecular subtypes were recently defined based on expression of master neurogenic transcription factors (TFs) *ASCL1* (SCLC-A) and *NEUROD1* (SCLC-N) and a rarer subtype defined by the non-neuroendocrine (Non-NE) Tuft Cell TF *POU2F3* (SCLC-P)^11,12^. SCLC expressing an immune signature without these TFs was defined as ‘inflamed’ (SCLC-I)^13^. Preclinical studies suggest subtype-dependent therapeutic vulnerabilities^14^ heralding potential for stratified therapy in clinical trials, potentially guided by ctDNA methylation subtyping^15^ where serial liquid biopsy could assess evolving subtype plasticity^16^.

Patients with SCLC have prevalent circulating tumour cells (CTCs)^17^, prompting our establishment of CTC-Derived patient eXplant (CDX) models in immunodeficient mice to explore SCLC biology and test novel therapeutics^12^. ASCL1 and/or NEUROD1 subtype CDX consist primarily of NE cells with a minority Non-NE subpopulation^12,18^ consistent with the NE to NonNE phenotype switch brought about by Notch signalling generating intra-tumoral heterogeneity^16,19^. POU2F3 expressing CDX13 tumours are exclusively Non-NE^12^. YAP1, initially considered a subtype determinant of SCLC^11^, is expressed in Non-NE cells within ASCL1 or NEUROD1 CDX^18^.

We recently described a subset of SCLC CDX lacking expression of *ASCL1* or *POU2F3,* that instead expressed the neurogenic, basic helix-loop-helix TF *ATOH1*, which could be co-expressed with *NEUROD1*^12^. *ATOH1* was expressed in 4 CDX models from 3/31 SCLC patients (9.6%). Two of these CDX were generated from the same patient pre- and post-treatment and maintained ATOH1 expression.

ATOH1 is homologue of *Drosophila melanogaster Atonal*, first identified in sensory organs of developing embryos^20^. In mouse models, Atoh1 (or Math1) is critical for development and differentiation of sensory cell types, including granule cells in the brain, sensory inner ear hair cells, Merkel cells in the skin, and secretory cells in the intestine^21–27^. Atoh1, like Ascl1, engages Notch signalling through lateral inhibition to avoid aberrant cellular differentiation in brain and intestine^24,28,29^. ATOH1 impact in cancer is context-dependent, described as a tumour suppressor in colorectal cancer and an oncogene in medulloblastoma^30,31^. Functional role(s) of ATOH1 in SCLC are unknown.

Here we explore transcriptional programmes and cellular functions(s) regulated by ATOH1 in SCLC. Although rare in our CDX biobank compared to SCLC-A, we identified ATOH1 in a subset of patients’ tumours and in additional Patient-Derived eXplants (PDX) models^32^. We show that in SCLC cell lines and/or CDX models, ATOH1 regulates neurogenesis, maintains cell survival *in vitro* and promotes tumour growth and liver metastasis *in vivo*. Our study adds to the emerging landscape of SCLC heterogeneity, highlighting potential for subtype-stratified approaches for improved treatment outcomes.

## RESULTS

### ATOH1, MYCL and chemosensitivity

We suggested ATOH1 as a SCLC subtype determinant after noting its expression in 4/38 CDX models that were distinct upon unsupervised clustering of whole transcriptomes^12^ (Figure 1A-i). Four ATOH1 CDX were derived from three donors: one sampled prior to chemotherapy (CDX25), one post-chemotherapy (CDX30P) and one where paired CDX were generated pre- and post-chemotherapy (CDX17, CDX17P), with maintained ATOH1 expression^12^ (Table S1). Whilst ATOH1 can be co-expressed with NEUROD1 (Figure 1A-i), we confirmed and extended Principal Component Analysis (PCA) of transcriptomic data from 39 CDX (including SCLC-A CDX31P^18^) that separated *ATOH1* models from *NEUROD1*-only models and from models expressing *ASCL1* or *POU2F3* (Figure 1A-ii). As ATOH1 is expressed in Merkel cells and most Merkel cell carcinomas (MCCs)^33^, we checked whether ATOH1 CDX were in fact derived from CTCs from mis-diagnosed MCC primary tumours. MCC is characterised by the presence of oncogenic Merkel cell polyoma virus (MCPyV) in 80% of cases^34^. We detected MCPyV sequences in MCC patient samples from a publicly available dataset (PRJNA775071) but not in any ATOH1 SCLC CDX (Figure S1A). Because a minority of MCC expresses neither ATOH1 nor MCPyV, we performed differential gene expression analysis (DGEA) of ATOH1 CDX compared to the entire CDX biobank and applied a Merkel cell-specific gene signature^35^ (Table S2), which was not significantly enriched in ATOH1 CDX (Figure S1B), further supporting that ATOH1 CDX do not have a Merkel cell origin.

**Figure 1.**
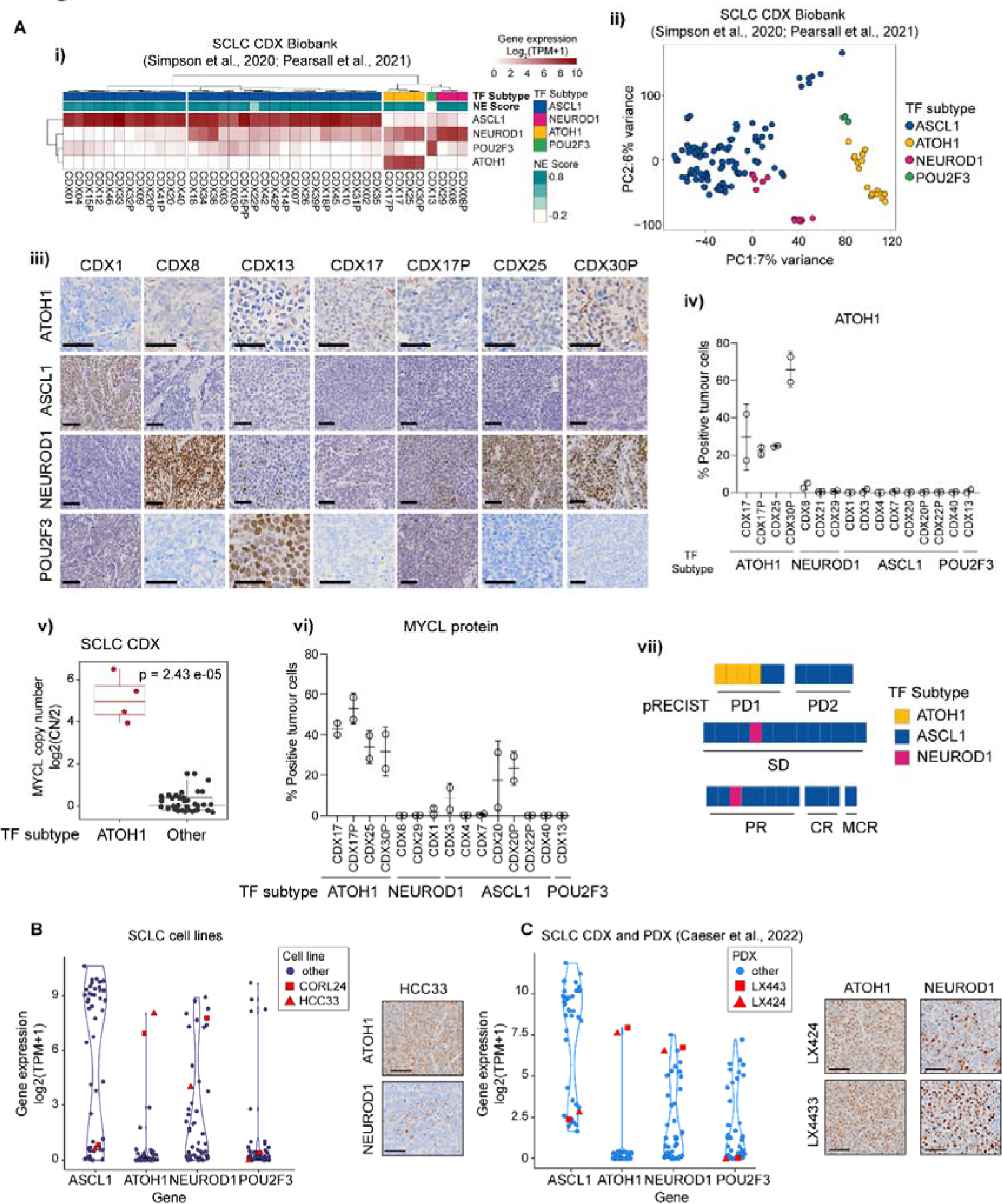
ATOH1 is expressed in a transcriptionally distinct subset of SCLC CDX, PDX and established cell lines. (A-i) Heatmap illustrating expression levels of *ASCL1, NEUROD1, ATOH1* and *POU2F3* in the SCLC CDX biobank, annotated by SCLC subtype and NE score^12,18^. Gene expression is shown as log_2_(TPM+1). (A-ii) Unbiased principal component analysis (PCA) of SCLC CDX in the biobank annotated by SCLC molecular subtypes. Key: blue, ASCL1; pink, NEUROD1; yellow, ATOH1; green, POU2F3. (A-iii) Representative IHC images for ATOH1, ASCL1, NEUROD1 and POU2F3 in a panel of CDX models belonging to different SCLC molecular subtypes. Scale bars: 50 μm. (A-iv) Quantification of ATOH1 expression in N=2 CDX tumours in a panel of CDX models. (A-v) Boxplot of MYCL copy number (CN), reported as CN ratio (Log2(CN/2)), in CDX grouped by molecular subtype (ATOH1 or other). Statistics reported as per Wilcoxon rank sum exact test. (A-vi) Quantification of MYCL expression by IHC in N=2 CDX tumours in a panel of CDX models belonging to different SCLC molecular subtypes (annotated below). (A-vii) Chemosensitivity scores of the SCLC CDX biobank according to pRECIST criteria, coloured by SCLC molecular subtypes. Key: yellow, ATOH1; blue, ASCL1; pink, NEUROD1. Data are reported after 1 cycle of cisplatin/etoposide treatment and as average of N>3 mice for N=29 CDX (see methods). Statistical analysis was performed with a Fisher’s exact test between ATOH1 CDX and the remaining CDX; p = 0.0049. (B-C) Violin plot representing expression of indicated NE and Non-NE TFs in SCLC established cell lines (B) and the SCLC CDX and PDX biobank^32^ (C); ATOH1-expressing HCC33, CORL24 (B) and LX424, LX443 (C) are highlighted in red. Gene expression is reported as Log_2_(TPM+1). Inserts are representative images of ATOH1 and NEUROD1 IHC staining for HCC33 (B) and LX424, LX443 (C).

SCLC subtyping was based predominantly on transcriptomes^11,13,36^. To examine ATOH1 protein expression we optimised an IHC assay using a commercially available antibody (from here on referred to as Ptech), that revealed nuclear ATOH1 staining only in ATOH1 subtype CDX (Figure 1A-iii, quantified in 1A-iv). Like ASCL1 and POU2F3 and in contrast to NEUROD1, ATOH1 transcript and protein expression followed a bimodal pattern; ATOH1 was either highly expressed or undetectable (Figure 1A-i, 1A-iii, 1A-iv). Whilst ATOH1 CDX expressed neither *ASCL1* nor *POU2F3* (Figure 1A-i), ATOH1 was expressed alone (CDX17P) or in combination with NEUROD1 at the transcript (Figure 1A-i) and protein level (Figure 1A-iii, CDX25, CDX30P: high *NEUROD1* expression, 78% positive tumour cells; CDX17: moderate *NEUROD1* expression, 30% positive tumour cells).

*MYCL* amplification is often observed in SCLC and MCC^37,38^. ATOH1 expression in CDX strongly correlates with *MYCL* focal amplification (Figure 1A-v, p=2.43*10^-5^), resulting in higher levels of *MYCL* transcript (Figure S1C) and MYCL protein (Figure 1A-vi, S1D) compared to other subtypes.

CDX reflect chemosensitivity profiles of their patient donors^12,39^. We investigated responses of ATOH1 CDX models to SoC (cisplatin/etoposide) *in vivo* adopting a modified version of preclinical RECIST (pRECIST) (see methods); tumour growth data are transformed to progressive disease (PD1, PD2), stable disease (SD) and partial (PR), complete (CR) and maintained responses (MCR)^40,41^. Compared to other molecular subtype CDX (31 SCLC-A, 25 patients, 2 SCLC-N, 2 patients) which displayed variable chemotherapy responses, all 4 ATOH1 CDX (3 patients) were the most chemoresistant, scoring as PD1 (Figure 1A-vii, Fisher’s exact test, p = 0.0049; Table S1). This finding was mirrored in clinical data from the 3 ATOH1 CDX patient donors who all had chemorefractory disease (Table S1). Whilst a larger number of ATOH1 models are required, our early findings imply a putative association of ATOH1 with chemotherapy resistance.

ATOH1 was expressed (transcript and protein) in 2/51 SCLC cell lines^42^ (Figure 1B) and 2/42 SCLC PDX^32^ (Figure 1C). The PDX and cell lines also exhibited bimodal ATOH1 expression accompanied by either low (HCC33) or high expression of NEUROD1 (CORL24, LX424, LX443) (Figure 1B-C, inserts). *MYCL* amplification was observed in ATOH1-expressing SCLC cell lines^43^ (HCC33 CN ratio ∼5, CORL24 CN ratio ∼2) and PDX (LX424/443^32^) and all ATOH1 preclinical models express amongst the highest reported levels of *MYCL* (Figure S1E-F). The ATOH1 expressing PDX were obtained from one chemorefractory donor (Table S1). Overall, whilst requiring larger sample sizes, these findings indicate that ATOH1 expression in SCLC CDX, PDX and cell lines, with or without NEUROD1, correlates with high *MYCL* expression and chemoresistance.

### ATOH1 in SCLC clinical specimens

*ATOH1* was detected in 1/81 SCLC tumours^36^ and in 3/100 small cell NE pulmonary and extrapulmonary carcinoma (SCNC) biopsies^44^. We detected *ATOH1* in 1/19 SCLC tumours profiled by single cell RNA-Seq (scRNA-Seq)^45^, previously classified as NEUROD1 subtype with expression of *NEUROD2* and *NEUROD4* (Figure 2A). We quantified ATOH1 protein in 65 specimens from 11 LS and 54 ES SCLC patients from the CHEMORES protocol and 37 specimens from LS patients enrolled in the CONVERT trial (methods, Table S4). ATOH1 was detected in 16/102 (16%) cases (Figure 2Ai-ii). One patient sample co-expressed ATOH1 and NEUROD1 (1/16, 6%) (Figure 2A-iii, Table S5) but in contrast to CDX and PDX, 8/16 (50%) ATOH1+ samples also had detectable ASCL1 expression and all three neurogenic TFs were detectable in 3/16 (19%) cases (Figure 2A-iii). Due to scant biopsies, we could not investigate cellular co-expression of TFs. ATOH1 expression did not correlate with altered OS or PFS compared to other SCLC subtypes (data not shown) in this cohort. Nevertheless, the relatively high prevalence of ATOH1 expression in clinical samples either alone or combined with ASCL1 and/or NEUROD1 encouraged further study of ATOH1-driven biology.

**Figure 2.**
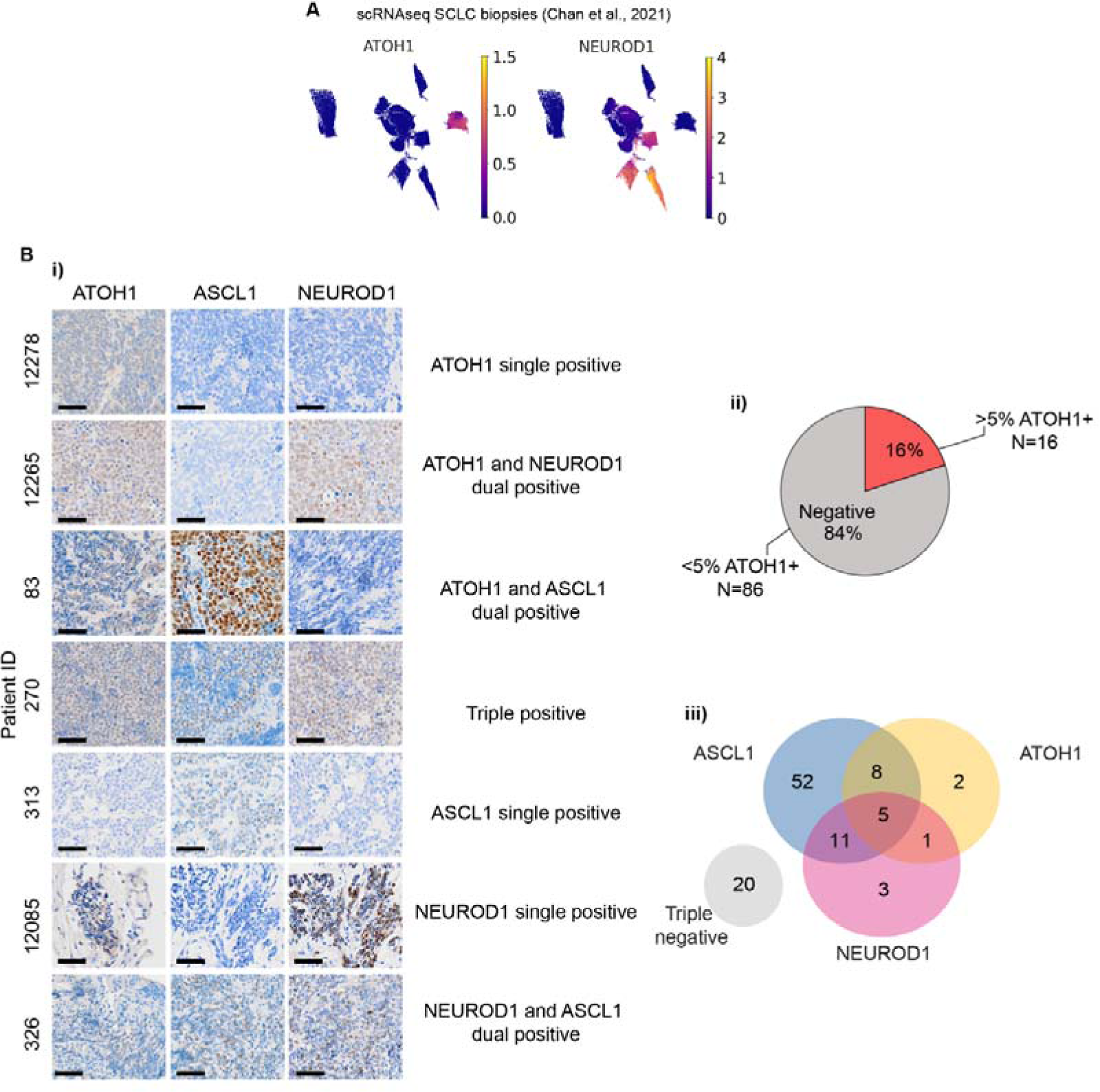
ATOH1 protein is expressed in SCLC clinical samples. (A) UMAP plots of single cell RNA-Seq (scRNA-Seq) from SCLC biopsies from the publicly available MSK SCLC Atlas^45^ reporting expression of *ATOH1* (left panel) and *NEUROD1* (right panel). Gene expression reported in units of log_2_(X + 1) where X = normalized counts. (B-i) Representative IHC images for ATOH1, ASCL1 and NEUROD1 in SCLC tissue biopsies presenting with single, dual or triple positivity (annotated). (B- ii) Pie chart illustrating the prevalence of ATOH1-positive (>5% positive tumour cells) clinical specimens (N=16/102). (B-iii) Venn diagram illustrating overlap of ASCL1, ATOH1 and NEUROD1 expression in N=102 clinical specimens as detected by IHC. Positivity determined as >1.5% positive tumour cells for ASCL1 and NEUROD1; positivity for ATOH1 determined as in B-ii.

### ATOH1 regulates a neurogenesis program by binding to E-boxes at promoter and enhancer regions in SCLC CDX

To interrogate the biological role of ATOH1 in CDX, we developed stable CDX17P lines carrying doxycycline-inducible (DOX) ATOH1 knock down (KD) ShRNA constructs (ShATOH1#1, -#3) or a control ShRNA targeting Renilla luciferase^46^ (ShRen) which also expressed GFP following DOX induction (Figure 3A-i). GFP expression enabled flow cytometric sorting of transduced cells. Maximal ATOH1 KD was observed after 7 days with both the Ptech antibody (Figure S2A) shown previously for IHC, as well as a previously in-house generated antibody (SY0287) (S2B-E, 3A-ii).

**Figure 3.**
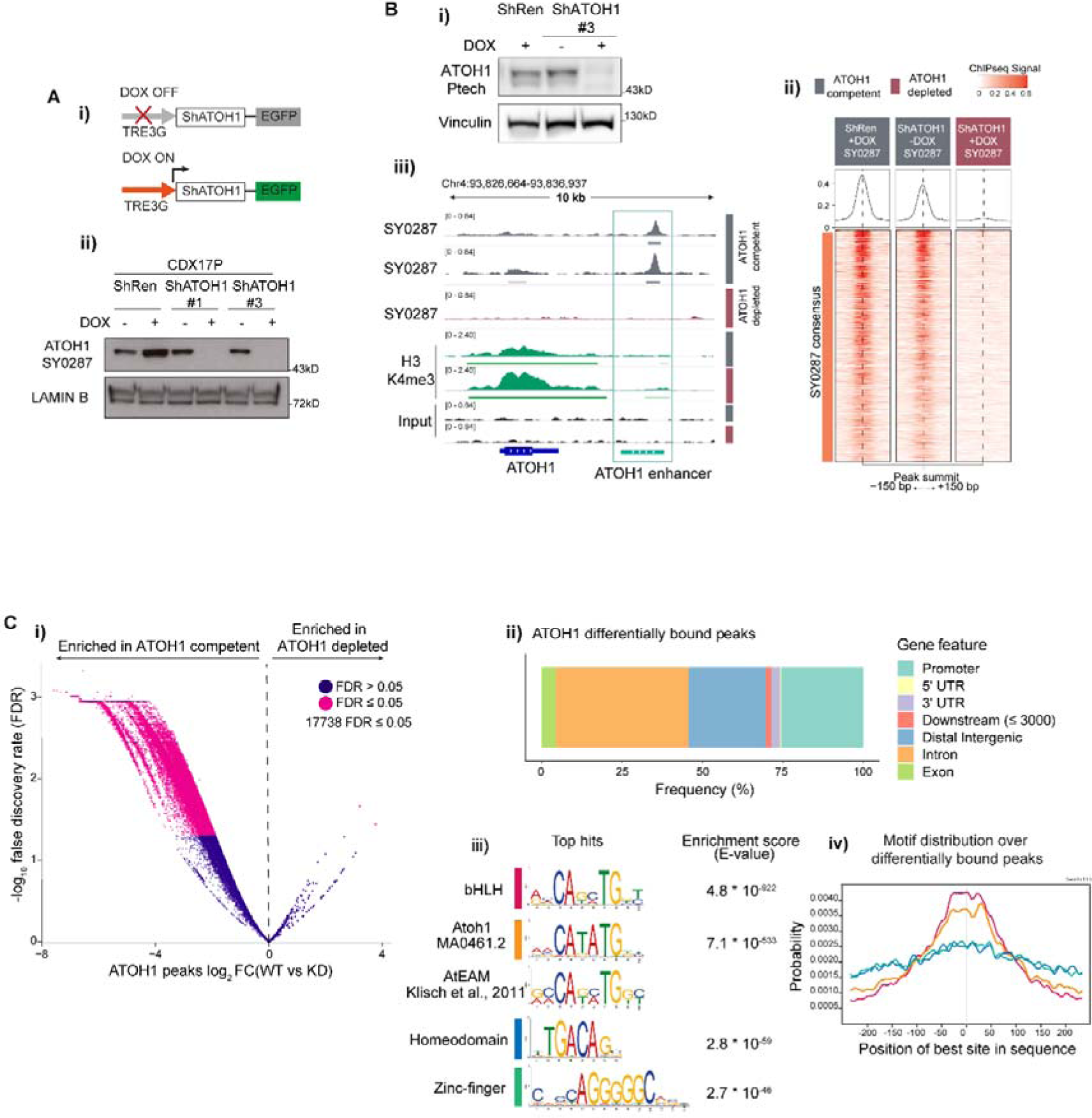
High confidence ATOH1 binding sites are located at promoter and enhancer regions and are enriched for E-box motifs. (A-i) Schematic of DOX- inducible knock-down (KD) system: without DOX, eGFP and shRNAs targeting ATOH1 (ShATOH1) or Renilla Luciferase (ShRen) are not expressed; upon induction with DOX, both eGFP and ShATOH1 or ShRen are expressed. (A-ii) (F) Nuclear fractionation validating ATOH1 KD with the in-house ATOH1 antibody SY0287 in CDX17P ShRen, ShATOH1#1 and ShATOH1#3 upon treatment with DOX for 7 days. (B-i) Western blot showing ATOH1 expression (detected with the Ptech antibody) in the samples processed for ChIP-Seq. (B-ii) Heatmap of ChIP-Seq signal for consensus peak sets SY0287 in ATOH1 competent (grey) and depleted (red) CDX17P, generated with the generateEnrichedHeatmap function within profileplyr v1.8.1^100^. (B-iii) ATOH1 binding peaks at ATOH1 locus highlighting ATOH1 binding peaks at ATOH1 downstream enhancer (light green), which are lost upon ATOH1 depletion. In dark green, ChIP-Seq tracks for H3K4me3 at the ATOH1 locus. The peaks were visualized with the Integrated Genomics Viewer genome browser. (C-i) Volcano plot of ATOH1 differentially bound regions (by false discovery rate, FDR < 0.05) in ATOH1 competent vs ATOH1 depleted CDX17P. Significant peaks highlighted in pink (17,738). (C-ii) Relative frequency of ATOH1 differentially bound peaks in regulatory genetic regions. (C-iii) Motif enrichment analysis of ATOH1 differentially bound peaks with MEME ChIP^101^. Mouse Atoh1 E-box-associated motif (AtEAM^49^) reported for comparison with Atoh1 DNA binding motif and bHLH motif. (C-iv) Centrimo^50^ analysis of the location of enriched motifs in ATOH1 differentially bound peaks.

Transcriptional programs of ATOH1 are unexplored in SCLC. To reveal ATOH1- specific TF-DNA binding we conducted chromatin immunoprecipitation with massively parallel sequencing (ChIP-Seq) on ATOH1-competent CDX17P (ShRen, 7 days DOX and untreated ShATOH1#3) and ATOH1-depleted ShATOH1#3 CDX17P (7 days DOX). Upon ATOH1 KD (Figure 3B-i), samples clustered based on ATOH1 expression (Figure S3A). Whilst ATOH1 ChIP-Seq signal was almost completely lost upon ATOH1 KD using SY0287 (Figure 3B-ii), some ChIP-Seq signal (∼50%) was retained with Ptech (Figure S3B) possibly due to non-specific antibody binding consistent with immunoblots (Figure S2A, 3B-i). Metagene analysis showed that ATOH1 peaks were located on the Transcription Start Site (TSS), near H3K4me3 peaks that identify active promoter regions^47^ and at intergenic regions mostly downstream of the gene body (Figure S3C) indicating that ATOH1 could regulate transcription at both promoter and distal regulatory elements. In support we found that ATOH1 binds to its own enhancer located downstream and highly conserved across species^22^ (Figure 3B-iii, S3D).

To identify high confidence ATOH1 binding peaks, we performed differential binding analysis between ATOH1 replete and depleted conditions, considering peaks detected by both antibodies and thus avoiding potential false positives. We found 17,738 ATOH1-specific binding events corresponding to 70% total peaks detected (25,464) (Figure 3C-i, Table S6). Amongst ATOH1-specific binding events, peaks are located at promoter regions (25%) and putative enhancer regions, such as distal intergenic (24%) and intronic regions (41%) (Figure 3C-ii) in accordance with recent results from MCC lines^48^. The most highly enriched motifs in ATOH1-specific peaks were basic helix-loop-helix binding motifs, including the reported ATOH1 DNA binding motif (MA0461.2) and the Atoh1 E-box-associated motif (AtEAM) identified in murine studies^22,49^ (Figure 3C-iii). Compared to the second and third most enriched motifs (homeodomains and zinc-fingers), E-box and ATOH1-specific motifs were found at the summit of ATOH1 peaks (Figure 3C-iv) suggesting they are uniquely present where there is highest ATOH1 signal^50^.

### ATOH1 target genes in SCLC CDX

We then sought to identify the biological processes in SCLC regulated by ATOH1 and its putative target genes. Consistent with its role as a neurogenic TF, ATOH1- bound genes were enriched in pathways related to neurogenesis (Figure S3E-F, Table S7). However, this analysis only considered DNA binding events irrespective of gene expression changes. To define genes directly regulated by ATOH1, we performed global transcriptomics (RNA-Seq) of CDX17P cells cultured *ex vivo* in presence or absence of DOX-induced ATOH1 KD (ShATOH#1, -#3). Genes directly regulated by ATOH1 should be downregulated after ATOH1 loss. As expected, ATOH1 was the most differentially expressed (DE) gene of ∼500 genes (Figure 4A-i, Table S8). Genes upregulated after ATOH1 KD included those involved in cell adhesion and migration, whereas downregulated genes play roles in neurogenesis (Figure 4A-ii, Table S9) and in inner ear hair cell differentiation, corroborated by decreased expression of independent inner ear hair cell signatures upon ATOH1 KD^51,52^ (Figure S4A-B, Table S10-S11). Overall, our findings agree with known ATOH1 transcriptional programs in murine developmental models whereby Atoh1 is required for inner ear hair cell and cerebellar granule cell development and differentiation^21^, although relevance of these processes to SCLC initiation and progression is unclear.

**Figure 4.**
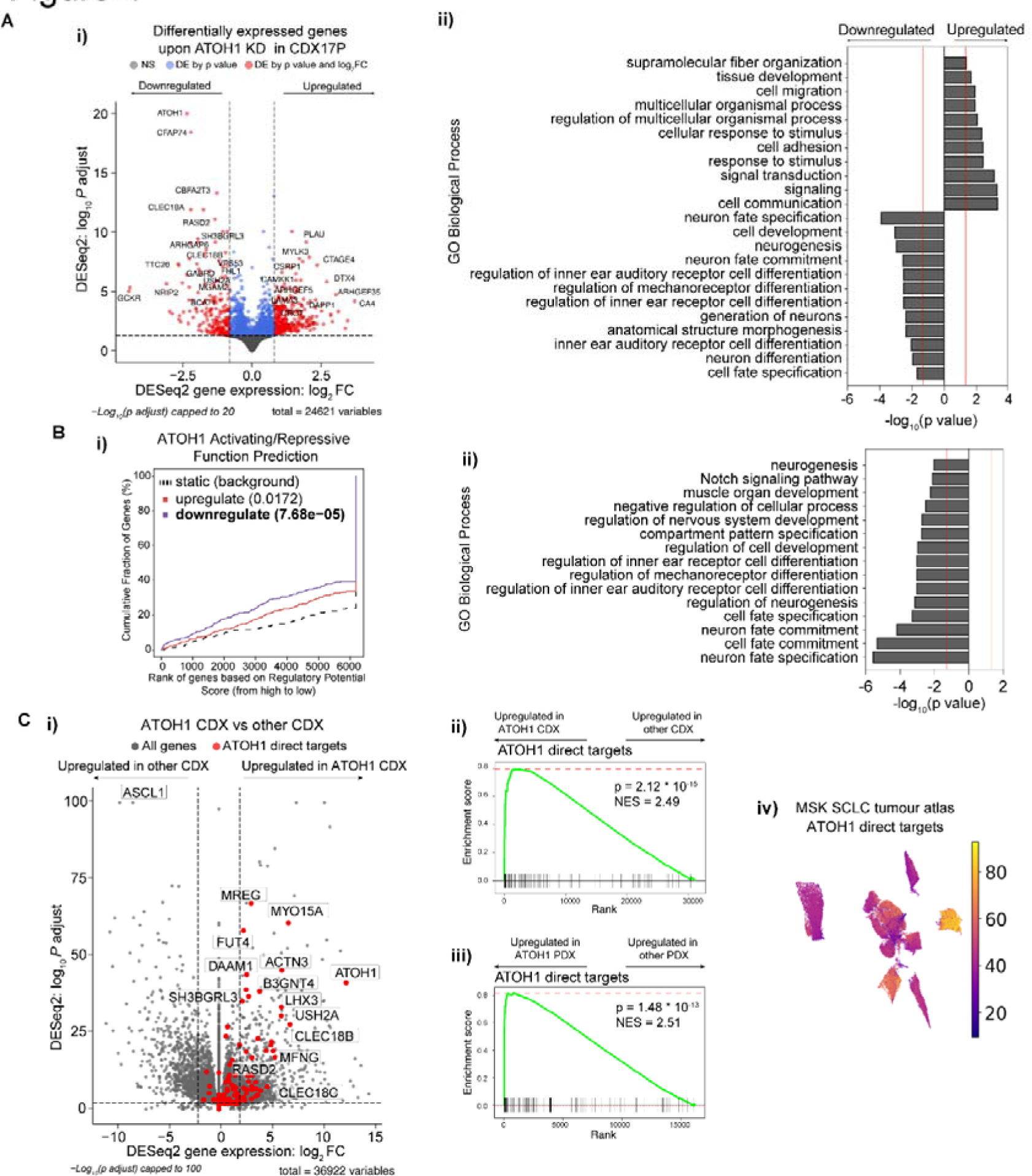
Identification of ATOH1 targetome and gene signature. (A-i) Volcano plot illustrating differentially expressed (DE) genes upon ATOH1 depletion (DOX treatment for 6 days) in CDX17P. Key: grey, not significant; blue, significant by p value; red, significant by p value <0.01 and log_2_(fold change) >0.8 or <-0.8. Dotted lines represent the thresholds for determining significant gene expression changes (p value <0.01 and log_2_(fold change) >0.8 or <-0.8). The most significant DE genes are labelled. (A-ii) Bar plot illustrating the top 20 biological processes up- and downregulated upon ATOH1 KD in CDX17P. Analysis was performed with gProfiler2^102^. (B-i) Prediction of ATOH1 transcriptional function after integration of ChIP-Seq and RNA-Seq with BETA^55^. ATOH1 KD results in downregulation of genes with ATOH1 binding sites identified in ChIP-Seq (p = 7.68 * 10^-5^) and is predicted to have a function in promoting transcription. (B-ii) Bar plot illustrating biological processes (performed with gProfiler2) associated with ATOH1 target genes identified in B-i. (C-i) Volcano plot illustrating genes enriched in ATOH1 CDX (N=4) compared to the whole CDX biobank (N=35). ATOH1 gene signature (i.e. ATOH1 target genes) highlighted in red. Dotted lines represent the thresholds for determining significant gene expression changes (p value <0.01 and log_2_(fold change) >2 or <-2). (C-ii) Gene set enrichment analysis (GSEA) for ATOH1 direct targets in ATOH1 CDX (N=4) vs the rest of the biobank (N=35). NES: normalised enrichment score. (C-iii) GSEA for ATOH1 direct targets in ATOH1 PDX (N=2) vs the rest of the MSK PDX biobank (N=40). GSEA analysis was performed with Fgsea^103^. (C-iv) UMAP of cumulative expression of ATOH1 direct targets in scRNA-Seq of SCLC tumour biopsies^45^. Expression of ATOH1 target genes is highest in the only ATOH1- expressing tumour (identified in Figure 2A).

ASCL1 and NEUROD1 are highly expressed in NE subtypes of SCLC^11,53^ and drive a NE transcriptional program. Given that ATOH1 also regulates neurogenesis, we asked whether NE status was affected by ATOH1 depletion. Whilst a 25-gene NE signature^54^ and SYP expression were unchanged upon ATOH1 KD (Figure S4C,E, Table S10), a 25-gene Non-NE signature was upregulated^54^ (Figure S4D, Table S10) suggesting that ATOH1 may contribute to NE to Non-NE plasticity, albeit without increased expression of YAP1 nor MYC (Figure S4E).

Fewer significant transcriptional changes were seen upon ATOH1 KD relative to the abundance of ATOH1 binding sites (by ChIP-Seq), suggesting that ATOH1 activity might be restricted to a subset of ATOH1-bound genes in SCLC CDX. Thus, to infer direct ATOH1 transcriptional targets in SCLC, we performed an integrated analysis of ChIP-Seq and RNA-Seq with the Binding and Expression Target Analysis (BETA)^55^. We found that ATOH1 mainly acts as a transcriptional activator (Figure 4B-i, blue line) and identified 150 genes downregulated upon ATOH1 depletion, directly downstream of ATOH1 (Table S12). Among these genes were components of Notch signalling (including *HES6, DLL1, DLL3, DLL4*) consistent with the interplay between ATOH1 and Notch signalling during brain and intestinal development^24,56^ and genes important for inner ear hair cell development such as *USH2A, LHX3* and *RASD2*^52^. Concordant with transcriptomics analysis (Figure 4A-ii), ATOH1 direct targets are also involved in neurogenesis and inner ear hair cell differentiation (Figure 4B-ii, Table S13).

This integrated analysis was performed in only CDX17P, so we next asked whether ATOH1 direct targets were conserved across all ATOH1 expressing CDX. We performed DGEA between ATOH1 CDX (CDX17, 17P, 25, 30P) and the whole CDX Biobank (35 CDX) (Figure 4C-i, Table S14), followed by gene set enrichment analysis (GSEA) for ATOH1 direct targets to demonstrate ATOH1 direct target genes were conserved (Figure 4C-ii, NES = 2.48, p = 1.13 * 10^-16^). We also detected high expression of ATOH1 target genes in the 2 ATOH1 SCLC PDX (Figure 4C-iii, NES = 2.44, p = 5 * 10^-10^) and an ATOH1 expressing tumour from the MSK SCLC tumour atlas dataset^45^ (Figure 4C-iv). These direct targets comprise the first SCLC-based ATOH1 gene signature consistently observed in CDX, PDX and tumour biopsies, indicative of a conserved transcriptional role for ATOH1 in SCLC.

### Impact of ATOH1 on SCLC CDX cell survival *ex vivo*

We examined the biological effects of ATOH1 depletion via DOX-inducible ATOH1 KD in CDX17P cells. Maximal ATOH1 KD was achieved after 7 days of DOX (Figure S2A) and was maintained for 14 days (the longest duration of *ex vivo* studies). Withdrawal of DOX restored ATOH1 expression (7 days +DOX, then 7 days -DOX) (Figure 5A-i, ii). ATOH1 depletion caused >50% decrease in cell viability (ShATOH1#1, p=0.0025; ShATOH1#3, p=0.0124), compared to un-induced and ShRen controls, which was attenuated by restoring ATOH1 expression (Figure 5B-i). To interrogate the mechanism of decreased cell viability, we established DOX-inducible ATOH1 KD in CDX30P and HCC33 SCLC cells (Figure S5A-B) and assessed cell death and cell cycle progression following ATOH1 depletion. Compared to ShRen DOX-induced controls and un-induced cells, there were no reproducible changes in cell cycle progression in CDX17P or CDX30P upon ATOH1 depletion for 14 days (Figure 5B-ii, Figure S5C). A modest ∼12% decrease in cell proliferation was evident in HCC33 cells although this did not constitute a complete proliferation arrest with ∼15% cells still cycling (Figure S5D). These slightly different effects on proliferation in CDX versus HCC33 may result from differences between cell lines and CDX *ex vivo* cultures. Instead, ATOH1 depletion increased cell death in CDX17P (55%), CDX30P (42%) and HCC33 (44%) after 14 days of ATOH1 depletion (Figure 5B-iii) via a caspase-3-independent process (Figure 5B-iv). After 7 days of DOX treatment, ATOH1 KD already induced detectable cell death (Figure 5C-i) and a decrease in ATP production, used as a proxy for viable cell number (Figure 5C-ii, iii, in red). Because other types of non-apoptotic, programmed cell death such as ferroptosis and pyroptosis have been observed in SCLC^57,58^, we induced ATOH1 KD in CDX17P and CDX30P ShATOH1#1 with DOX, with or without cell death pathway inhibitors for 7 days. Inhibition of apoptosis, pyroptosis, necroptosis or ferroptosis (with single or combined inhibitors) did not prevent ATOH1 KD-induced loss of cell viability (Figure 5C-ii, iii). Taken together, these findings identify ATOH1 as necessary for cell survival in CDX17P, CDX30P and HCC33 cells as its depletion induces cell death, either via an undefined programmed cell death pathway or most likely via necrosis.

**Figure 5.**
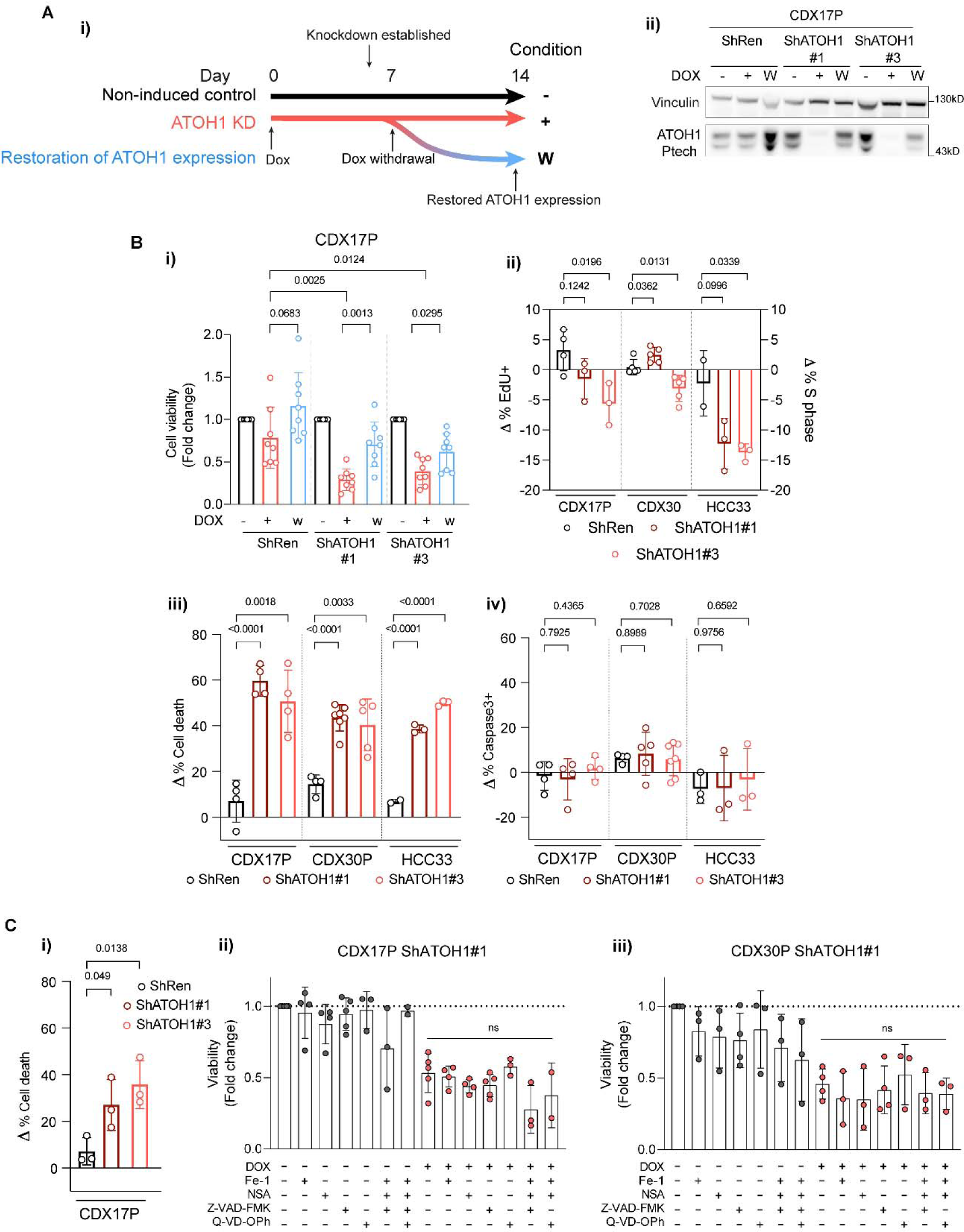
ATOH1 is necessary for SCLC cell survival *in vitro*. (A-i) Schematic of induction of ATOH1 KD. ATOH1 KD was established after 7 days induction with 1 μg/ml doxycycline (DOX). Cells were cultured for a total of 14 days with DOX (red line, +) or without DOX as controls; after the initial 7 days induction with DOX, a part of cells was plated without DOX to restore ATOH1 expression (blue line, W). Untreated parental cells served as additional control (black line, -). (A-ii) Western blot validation of ATOH1 depletion and restoration in the conditions specified in A-i. ShRen treated with DOX for 14 days and untreated ShRen, ShATOH1#1, ShATOH1#3 and were used as control. (B-i) Relative cell viability measured with CellTiter-Glo® (Promega) upon ATOH1 KD (red) and restoration (blue) compared to un-induced controls (black). N=8 independent experiments. (B-ii) Flow cytometry quantification of cell cycle progression by EdU (CDX17P, HCC33) and PI incorporation (CDX30P). Data was normalised to DOX-untreated parental controls by subtracting the proportion of cells in S phase in untreated cells to that of DOX-treated cells (Δ % S phase = % S phase_DOX-treated_ - % S phase_untreated_); ShATOH1 conditions were then compared to ShRen controls. CDX17P, N=4 ShRen, N=3 ShATOH1#1 and #3; CDX30P, N=5; HCC33, N=2 ShRen, N=3 ShATOH1#1 and #3 independent experiments. (B-iii) Flow cytometry quantification of cell death after 14 days induction with DOX of ATOH1 KD, normalised as in B-ii. Total cell death is reported as sum of apoptotic and necrotic cells. CDX17P: N=4; CDX30P: N=4 ShRen, N=7 ShATOH1#1, N=5 ShATOH1#3; HCC33: N=2 ShRen, N=3 ShATOH1#1 and #3 independent experiments. (B-iv) Same as B-iii, reporting total Caspase-3 positive cells. All statistics in panel B are reported as two-tailed unpaired *t* tests across indicated conditions. C-i) Flow cytometry quantification of cell death (as defined in B-iii) after 7 days DOX-induction of ATOH1 KD in CDX17P. N=3 independent experiments. P values are reported in panel B and C-i as per two-tailed unpaired *t* test. (C-ii, C-iii) ShATOH1#1 CDX17P (C-ii) and CDX30P (C-iii) cells were treated with (red) or without (black) DOX and with or without ferrostatin-1 (1µM), necrosulfonamide (NSA, 100 nM) or Z-VAD-FMK/Q-VD-OPh (20µM) and indicated combinations for 7 days. Cell viability was measured with CellTiter-Glo®, normalized to vehicle treated, DOX-untreated cells and reported as fold change. Statistics in C-ii and C-iii are reported as per one-way ANOVA test with Dunnett’s test correction for multiple comparisons between DOX-treated conditions with and without programmed cell death inhibitors. Data are shown as mean ± SD.

### Impact of ATOH1 on tumour growth *in vivo*

We next asked whether the role of ATOH1 in maintaining cell survival *ex vivo* translated to an impact on tumour growth *in vivo*. CDX17P control ShRen or ShATOH1(#3) cells were implanted subcutaneously (s.c.) in immunocompromised mice, and KD was induced with DOX-supplemented feed after 19 days (Figure 6A), when mice had palpable tumours. Once tumours reached 500-800 mm^3^ they were surgically resected and mice kept on study for 28 days to allow time for metastatic dissemination (based on previous experiments, see methods, Figure 6A).

**Figure 6.**
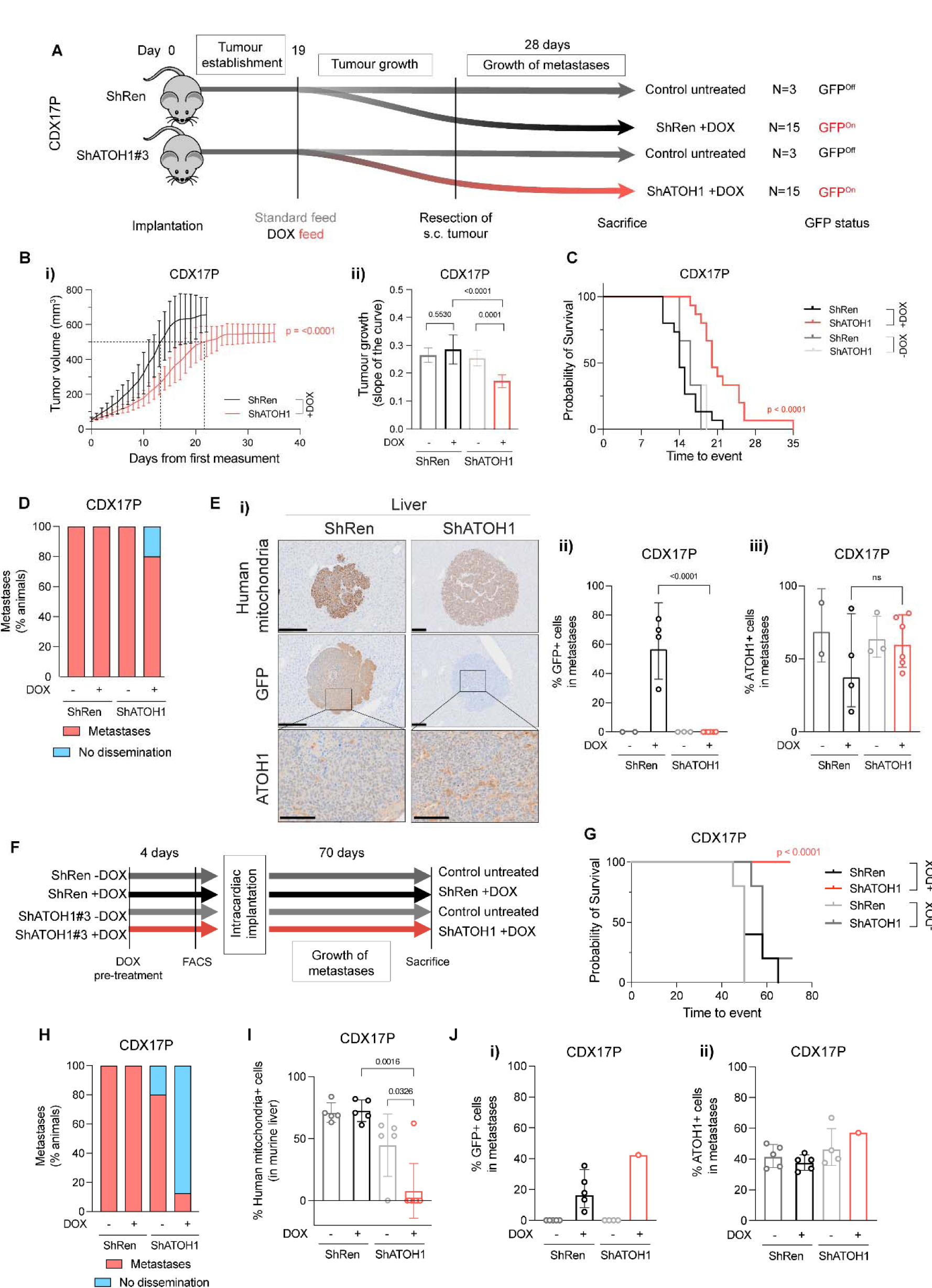
ATOH1 depletion decreases tumour growth kinetics and metastasis *in vivo*. (A) *In vivo* study design to investigate subcutaneous (s.c.) tumour growth and metastasis after s.c. tumour resection. CDX17P ShRen and ShATOH1#3 (ShATOH1) were injected s.c. in NSG mice and left for 19 days to allow for tumour establishment. After 19 days, mice were fed either standard diet (control arms, N=3) or DOX-supplemented feed (experimental arms, N=15) and s.c. tumour growth was assessed. S.c. tumours were surgically resected when at 500-800 mm^3^ to allow for metastatic dissemination and mice were kept on study for 28 days or until s.c. tumour reached maximum size, whichever came first. (B-i) S.c. tumour growth curves, from day of first tumour measurement to s.c. tumour resection (see methods), of mice implanted with ShRen and ShATOH1 and fed DOX-supplemented diet. Key: black, ShRen fed DOX-diet; red, ShATOH1#3 fed DOX-diet. N=15 mice per cohort; data reported as mean ± SD. Dotted lines indicate when tumours from each cohort reached 500 mm^3^: ShRen, 14 ± 3 days; ShATOH1, 21 ± 5 days. (B-ii) Quantification of the slope of tumour growth curves in B. Key: same as in B; shades of grey for control cohort fed standard diet for the duration of the study. P values were calculated with ANCOVA test and slope of the curve was reported as mean ± SD for each cohort. (C) Kaplan-Meier curve of time to surgical resection of s.c. tumour or maximum 800 mm^3^ for inoperable tumours. Control arms, fed a standard diet, reported in scales of grey. P values were calculated with Log-rank Mantel-Cox test. (D) Quantification of metastatic dissemination to the liver in N=3 mice fed standard diet, N=5 ShRen- and N=15 ShATOH1-tumour bearing mice fed DOX-diet that underwent surgical resection of s.c. tumour and survived on study for at least 22 days after resection. Data is shown as percentage of animals displaying metastatic dissemination (disseminated tumour cells and micro/macro-metastases, in red) or no metastatic dissemination in the liver (blue). Metastases were identified based on human mitochondria staining. (E-i) Representative images of human mitochondria, GFP and ATOH1 IHC staining in liver from ShRen DOX-fed and ShATOH1#3 DOX-fed cohort. Scale bars: 200 μm for human mitochondria and GFP; 100 μm for ATOH1. (E-ii, E-iii) Quantification of GFP (E-ii) and ATOH1 (E-iii) IHC staining in metastases from N=2 DOX-untreated ShRen, N=3 DOX-untreated ShATOH1#3, N=4 ShRen DOX-fed, N=6 ShATOH1#3 DOX-fed mice. Data are shown as geometric mean ± geometric SD. P values are reported as per two-tailed unpaired Mann Whitney U test. (F) *In vivo* study design to investigate development of metastasis following intracardiac implantation. Prior to cell implantation, ATOH1 depletion was induced by DOX treatment for 4 days *in vitro*, followed by sorting GFP- positive, viable cells by flow cytometry. Untreated control cells were sorted exclusively for viable cells. Animals in the DOX treatment cohorts were fed a DOX-supplemented diet 24 hours prior to implantation and they were kept on that diet until endpoint. Animals in the uninduced control groups were given a standard diet. Animals from all 4 cohorts (ShRen +/- DOX and ShATOH1 +/- DOX) were removed at the onset of symptoms (i.e., distended abdomen, detailed in methods) or after 70 days. (G) Kaplan-Meier curve of time to sacrifice. Control cohorts, fed a standard diet, reported in scales of grey. P values were calculated with Log-rank Mantel-Cox test. (H) Quantification of metastatic dissemination to the liver for each cohort. Data is shown as per Figure 6D. (I) Quantification of metastatic cells in the liver for each cohort. Metastatic cells were identified based on human mitochondria staining. Data shown as mean ± SD. P values were calculated with a two-tailed unpaired Mann Whitney U test. (J) Quantification of GFP (J-i) and ATOH1 (J-ii) IHC staining in metastases from N=5 DOX-untreated ShRen, N=5 DOX-untreated ShATOH1, N=5 ShRen DOX-fed, N=1 ShATOH1#3 DOX-fed mice. Data are shown as geometric mean ± geometric SD. No statistical test could be performed as ShATOH1 contained only one value.

A significantly delayed s.c. tumour growth was observed in mice bearing DOX- induced ATOH1 KD tumours compared to DOX-induced ShRen controls or un- induced tumours (Figure 6B-i, ii). This tumour growth delay extended time to reach the experimental endpoint tumour volume or s.c. tumour surgical resection (22 days for ShRen, 35 days for ShATOH1, p<0.0001, Figure 6C). To interpret the observed growth delay, we examined persistence of ATOH1 KD throughout the experiment by performing IHC for ATOH1 and GFP in resected s.c. tumours (mean tumour volume and time from implant: 603±54 mm^3^, 44±5 days ShRen +DOX; 552±48 mm^3^, 70±13 days ShATOH1 +DOX) (Figure 6B-i). At tumour resection, mice bearing DOX- induced ATOH1 KD tumours showed a 75% reduction in ATOH1 protein expression and both DOX-induced controls and KD tumours had high expression of GFP (Figure S6A-i, ii). However, GFP expression was ∼10% lower in DOX-induced ATOH1 KD tumours (Figure S6A-ii, p=0.008) and expression of GFP and ATOH1 was heterogeneous in DOX-induced ATOH1 KD tumours, with most tumour presenting with some GFP-, ATOH1+ regions (Figure S6A-iii).

Overall, these data indicate that reduced ATOH1 expression promotes tumour growth delay *in vivo,* where impact may have been attenuated by outgrowth of ATOH1 positive cells which are potentially un-transduced wild-type cells or cells that escaped inducible KD, as reported in other settings^59,60^. These data are consistent with a selective pressure to re-instate ATOH1 expression in ATOH1 KD tumours supporting a pro-tumorigenic role for ATOH1.

### A Role for ATOH1 in liver-metastatic dissemination *in vivo*

We previously reported metastasis to multiple organs, including brain and liver, occurs after resection of s.c. CDX17P tumours^12^. To investigate whether ATOH1 supports metastatic growth, s.c. tumours were resected and mice left on study for 28 days (Figure 6A) before metastasis (defined as >50 tumour cells) were quantified using a human mitochondria antibody and IHC. Dissemination, predominantly to the liver, was observed in all cohorts regardless of DOX feed, including single tumour cells, micro-or macro-metastasis (Figure 6D). Although frequency of liver metastases between control and DOX-induced ATOH1 KD mice was approximately equivalent, all liver metastases from DOX-induced ShATOH1 mice were negative for GFP and expressed similar levels of ATOH1 compared to un-induced tumours (Figure 6E-i, ii), again implying a selective pressure to retain/re-express ATOH1^59,60^ and indirectly suggesting a role for ATOH1 in promoting liver metastasis.

In a more direct approach to investigate the role of ATOH1 in metastasis, we performed intracardiac injection of tumour cells (Figure 6F), reasoning liver metastasis would occur faster, allowing less time for outgrowth of cells with high or re-expressed ATOH1 (Figure 6E). CDX17P control ShRen or ShATOH1 cells were cultured with or without DOX for 4 days to induce ATOH1 KD *in vitro* and GFP- positive viable cells were sorted by flow cytometry before intra-cardiac injection. One group of mice per construct (ShRen and ShATOH1) received DOX-supplemented feed (N=5 ShRen and N=8 ShATOH1), while control animals were maintained on standard diet (N=5 ShRen and N=5 ShATOH1). Animals were removed from study 70 days after intracardiac injection (see methods, Figure 6F).

Almost all animals (14/15) in control cohorts (standard feed or implanted with DOX- induced ShRen cells) were removed before study endpoint due to extensive metastatic liver disease (Figure S6B). In contrast, 8/8 (100%) animals implanted with DOX-induced ShATOH1 cells reached study endpoint (time from implantation: 53.6 ± 7.9 ShRen+DOX; 70 ± 0 ShATOH1+DOX; Figure 6G). There was a significant reduction in metastatic burden in animals with ATOH1 KD compared to control cohorts (Figure 6H-I) and only one animal in the DOX-induced ShATOH1 group developed liver metastasis (Figure S6B). Despite showing positive GFP expression (>40% GFP+ cells), the only liver metastasis derived from ATOH1 KD cells also exhibited ATOH1 positivity in >60% of metastatic cells, indicating that ATOH1 KD was not completely retained in these cells (Figure 6Ji-ii). These data provide more direct evidence that ATOH1 KD reduced metastasis to the liver and promoted longer survival.

## DISCUSSION

Emerging understanding of SCLC subtypes and phenotypic plasticity are considered key to support rational development of biomarker-directed personalised treatments^14^. Building upon knowledge of inter- and intra-tumoural heterogeneity^32,44^, we have characterised the ATOH1 subtype, defining its prevalence and demonstrating pro- tumour functions of growth and metastasis.

ASCL1, NEUROD1 and ATOH1 are all proneural TFs negatively regulated by Notch signalling^24,28,61^. Whilst expression of ATOH1 is not reported during normal lung development, its expression has been reported in NE lung cancer^62^, extrapulmonary high-grade neuroendocrine cancers^44^, Merkel cell carcinoma (MCC)^33^, medulloblastoma^63,64^ and rarely in NSCLC^65^ and colorectal cancer (CRC)^30,66,67^. Whilst mechanistically understudied, in medulloblastoma and MCC ATOH1 is tumour-promoting^31,68–70^, whereas it is a tumour suppressor in CRC^30,66^. These opposing context-dependent functions have been attributed to imbalance between differentiation and proliferation driven by abnormal ATOH1 expression levels^71^.

Co-expression of subtype TFs is commonly observed, contributing to SCLC heterogeneity^12,32,72,73^. ATOH1 was found to be frequently expressed in SCLC clinical samples, either alone or with ASCL1 and/or NEUROD1 (Figures 1, 2) extending existing sparse data^62^. In CDX30 where ATOH1 was co-expressed with NEUROD1, ATOH1 depletion impacted cell survival *ex vivo* (Figure 5), suggesting that NEUROD1 could not compensate for ATOH1 loss. Furthermore, NEUROD1 was not identified amongst ATOH1 direct targets and there was minimal overlap with ASCL1 and NEUROD1 target genes (Figure 4, Table S15). Like NEUROD1 and ASCL1 in their respective subtypes^74–79^, ATOH1 supports cell viability in ATOH1 subtype tumour cells (Figure 5).

In SCLC, ATOH1 exerts its function by binding E-box motifs at promoter and enhancer regions of target genes as in the developing mouse brain^49^ and in MCC^80^, including binding to its own downstream enhancer^22^ (Figure 3). In CDX, ATOH1 directly regulates expression of genes involved in neuronal fate development and mechanoreceptor differentiation (Figure 4) consistent with murine developmental studies^21,81,82^. This is also consistent with the role of ATOH1 in MCC^33^. The ability of ATOH1 to regulate neuronal fate determination and Notch ligands (DLL1, DLL3, DLL4) in mice^24^ mirrors the activity of ASCL1 in SCLC^53,74^; in CDX17P, ATOH1 depletion increased expression of Non-NE and cell adhesion genes invoking a similar role for ATOH1 in NE fate determination in SCLC (Figure S4). However, as the NE gene expression signature was retained upon ATOH1 depletion (Figure S4), additional factors, for example, MYC overexpression^16^, are likely required to promote full NE to Non-NE transition in ATOH1-driven SCLC. The need for additional signals to fully induce a NE to Non-NE transition is similarly posited in studies of ASCL1 and NEUROD1 depletion in SCLC, where morphological changes or a NE to Non-NE transition were not observed^77,78,83,84^.

Both ATOH1 and ASCL1 correlate with *MYCL* overexpression (Figure 1). In SCLC, overexpression/genetic amplification of *MYCL* was often correlated with the SCLC-A subtype and *MYCL* is a direct transcriptional target of ASCL1^35, 52, 86^. A more complex relationship was recently revealed by a clinical study whereby MYCL protein was present in only ∼30% of ASCL1+ samples^73^. Further adding to this heterogeneity, we show that all ATOH1-expressing CDX present focal amplification and overexpression of MYCL (Figure 1, S1). A correlation between ATOH1 and MYCL expression was also observed in MCC^37,38^. However, we did not identify MYCL as a direct ATOH1 target (Table S12) and *MYCL* expression was unchanged upon ATOH1 depletion (Table S8, Figure 4). Combined, these data indicate that other factors contribute to *MYCL* expression in ATOH1-positive SCLC.

The profound impact of metastasis on SCLC patient outcomes drives a pressing need to understand and target underlying mechanisms. Acquisition of neuronal gene expression programmes is associated with invasive and metastatic SCLC in cell lines and GEMMs^59,85,86^. In CDX17P, ATOH1 is pro-metastatic (Figure 6) drawing parallels with the ATOH1 pro-invasive phenotype in MCC^87^ and its pro-metastatic role in medulloblastoma^88^. ATOH1 downregulation was linked with loss of cell adhesion (Figure 4A-ii, Table S8), which was also observed in MCC^33,89^.

SCLC was once considered to derive from pulmonary neuroendocrine cell (PNEC) precursors^90^. However, elegant studies in SCLC GEMMs describe different potential cells of origin^59,91–93^ with differences only evident at the molecular level^16,45,53^. In this regard, similarities between MCC and ATOH1-driven SCLC are intriguing. MCC is a NE skin carcinoma, expressing epithelial and NE markers with morphological, ultrastructural and immunohistochemical features shared with Merkel cells^91–93^ yet there is no direct histo-genetic link between Merkel cells and MCC with ongoing debate on cell(s) of origin of MCC^94,95^. Tumour heterogeneity in MCC is attributed to variant disease aetiologies mediated by either UV exposure or Merkel cell polyomavirus (MCPyV) integration^95^. Virus-positive MCC has low mutation burden, whilst virus-negative MCC, like SCLC, have characteristic RB1 and TP53 mutations in a highly mutated landscape^96,97^. The recent identification of ‘mesenchymal-like’ MCC with an ‘inflamed’ phenotype exhibiting better response to immunotherapy draws parallels with the SCLC-I subtype^13^ and contrasts ‘immune-cold’ immunotherapy resistant MCC with higher expression of neuroepithelial markers including ATOH1^98^. Altogether, that the ATOH1 subtype of SCLC CDX shares features with NE SCLC and with MCC, another NE cancer, is perhaps not surprising and might indicate convergent tumour evolution^94,99^.

In summary, here we validate the ATOH1 SCLC subtype (SCLC-AT) where ATOH1 suppresses cell death and promotes tumour growth and metastasis. Further studies are now needed to deepen our understanding of ATOH1-driven SCLC biology and to address whether there are therapeutic vulnerabilities of this subtype.

## Acknowledgements

This work was supported through Core Funding to Cancer Research UK (CRUK) Manchester Institute (grant number A27412), Manchester CRUK Centre Award (grant number A25254), the CRUK Lung Cancer Centre of Excellence (grant number A20465), Cancer Research UK Manchester Centre award (CTRQQR-2021\100010), The Christie Charitable Fund, National Cancer Institute R35 CA263816 and U24 CA213274. Patient recruitment was supported by the National Institute for Healthcare Research (NIHR) Manchester Biomedical Research Centre, the NIHR Manchester Clinical research Facility at The Christie Hospital and the CRUK Lung Cancer Centre of Excellence. Sample collection was undertaken through the molecular mechanisms underlying chemotherapy resistance, therapeutic escape, efficacy, and toxicity improving knowledge of treatment resistance in patients with lung cancer or CHEMORES protocol, the TARGET (tumour characterization to guide experimental targeted therapy) study and the CONVERT protocol (concurrent once- daily versus twice-daily radiotherapy: a 2-arm randomised controlled trial of concurrent chemo-radiotherapy comparing twice-daily and once-daily radiotherapy schedules in patients with limited stage small cell lung cancer (SCLC) and good performance status). Dr. Frese, Dr. Simpson and Prof. Dive supervised and devised the study. Dr. Catozzi, Dr. Peiris-Pagès, Dr. Simpson, and Prof. Dive co-wrote the manuscript. Dr. Catozzi, Dr. Peiris-Pagès, Ms. Davies-Williams, Mr. Revill, and Mr. Morgan performed immunohistochemistry analysis, data analysis and interpretation. Dr. Catozzi carried out all experiments on CDX and cell lines, ChIP-Seq, RNA-Seq and western blotting, including data analysis and interpretation. Dr. Catozzi, Dr. Humphrey, Mr. Chen carried out bioinformatics analyses. Dr. Peiris-Pagès designed the *in vivo* metastases studies and analysed the metastatic dissemination of ATOH1 KD cells *in vivo*. Ms. Galvin, Mr. Roebuck and Dr. Lallo were responsible for all *in vivo* work described. Dr. Frese had oversight of all patients with circulating tumour cell–derived explant models and model generation and helped edit the manuscript. Dr. Pearce and Dr. Kerr had oversight of all bioinformatics analyses. Ms. Priest, Dr. Foy, Mr. Carter, and Prof. Blackhall oversaw the acquisition of ethical permission and patient consent and the collection of blood samples from patients in the CHEMORES study. Dr Rudin provided PDX and assisted with manuscript revision. Prof. Blackhall assisted with manuscript revision and is the chief investigator of the CHEMORES study. All authors read and approved the final manuscript.

---

**Figure S1.**
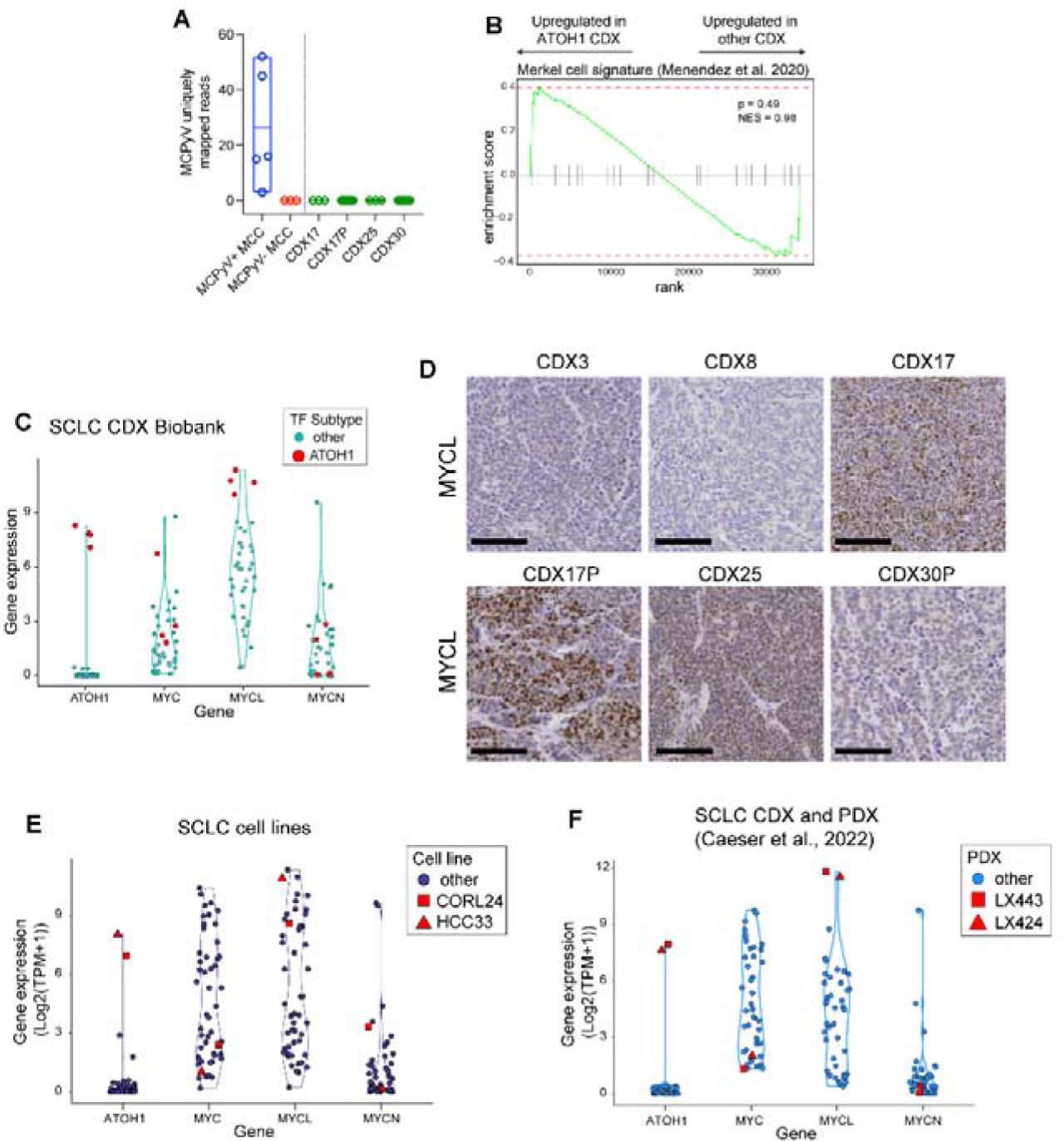
ATOH1 CDX do not have MCC origin and present high expression of MYCL. Relative to. Figure 1. (A) Detection of Merkel cell polyoma virus (MCPyV) transcripts in positive and negative control human Merkel cell carcinoma (MCC) samples (PRJNA775071) and ATOH1 CDX. (B) Gene set enrichment analysis (GSEA) for a Merkel cell gene signature from Menendez et al.^35^ in ATOH1 CDX (N=4) compared to the whole biobank (N=35). GSEA was performed with Fgsea^103^. (C) Violin plot of expression of indicated *MYC* family genes in the SCLC CDX biobank (N=39). ATOH1 subtype samples and preclinical models highlighted in red. (D) Representative IHC images for MYCL in SCLC-A CDX3, SCLC-N CDX8 and ATOH1 CDX CDX17, 17P, 25 and 30P. (E-F) Violin plot of expression of indicated *MYC* family genes in SCLC cell lines^42^ (E) and SCLC PDX^32^ (F) from publicly available datasets. ATOH1 subtype preclinical models highlighted in red and annotated by shape as in legend.

**Figure S2.**
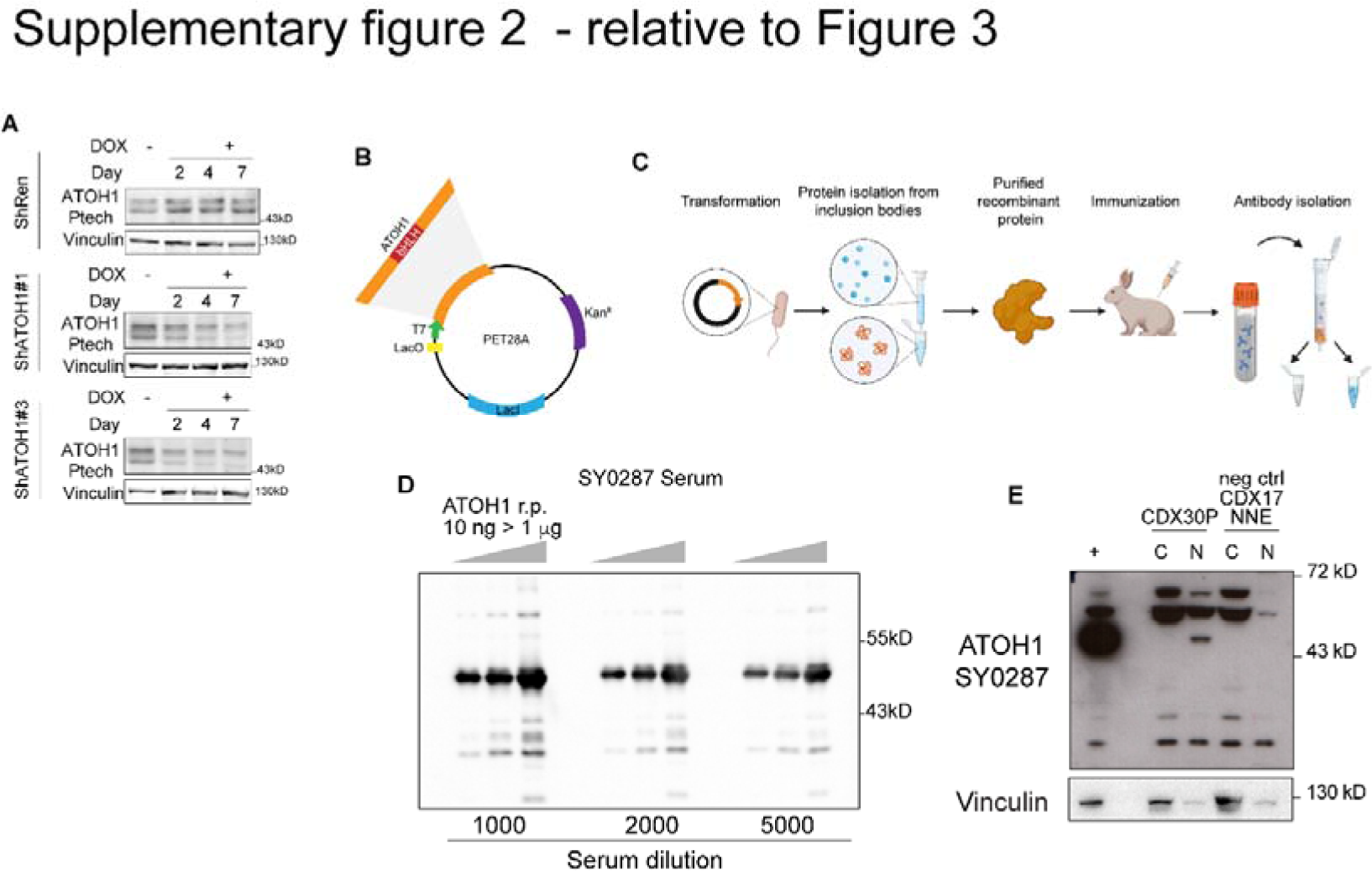
ATOH1 antibody production and validation. Relative to. Figure 2. (A) Western blot showing ATOH1 expression detected by the Ptech antibody over a time-course (0 to 7 days) of ATOH1 knockdown (KD) induction with doxycycline (DOX) in CDX17P. ShRen served as control for ATOH1 KD and Vinculin served as loading control. Western blots are representative of N=2 independent experiments. (B) Schematic of plasmid construct to express ATOH1 recombinant protein in IPTG- inducible PET28A system. (C) Workflow to produce the in-house antibody: ATOH1 recombinant protein was purified from bacterial culture and used for immunization of one rabbit. Polyclonal antibodies were isolated from final bleed serum by affinity purification. (D) Test of SY0287 serum before affinity purification against increasing amounts of ATOH1 recombinant protein (10 ng, 100 ng and 1 μg) by western blot. (E) Validation of ATOH1 detection by nuclear (N) and cytoplasmic (C) fractionation of CDX30P (positive control) and CDX17 Non-NE cells (Negative control). Transient ATOH1 overexpression in LentiX 293T cells (indicated as +) served as positive control for detection.

**Figure S3.**
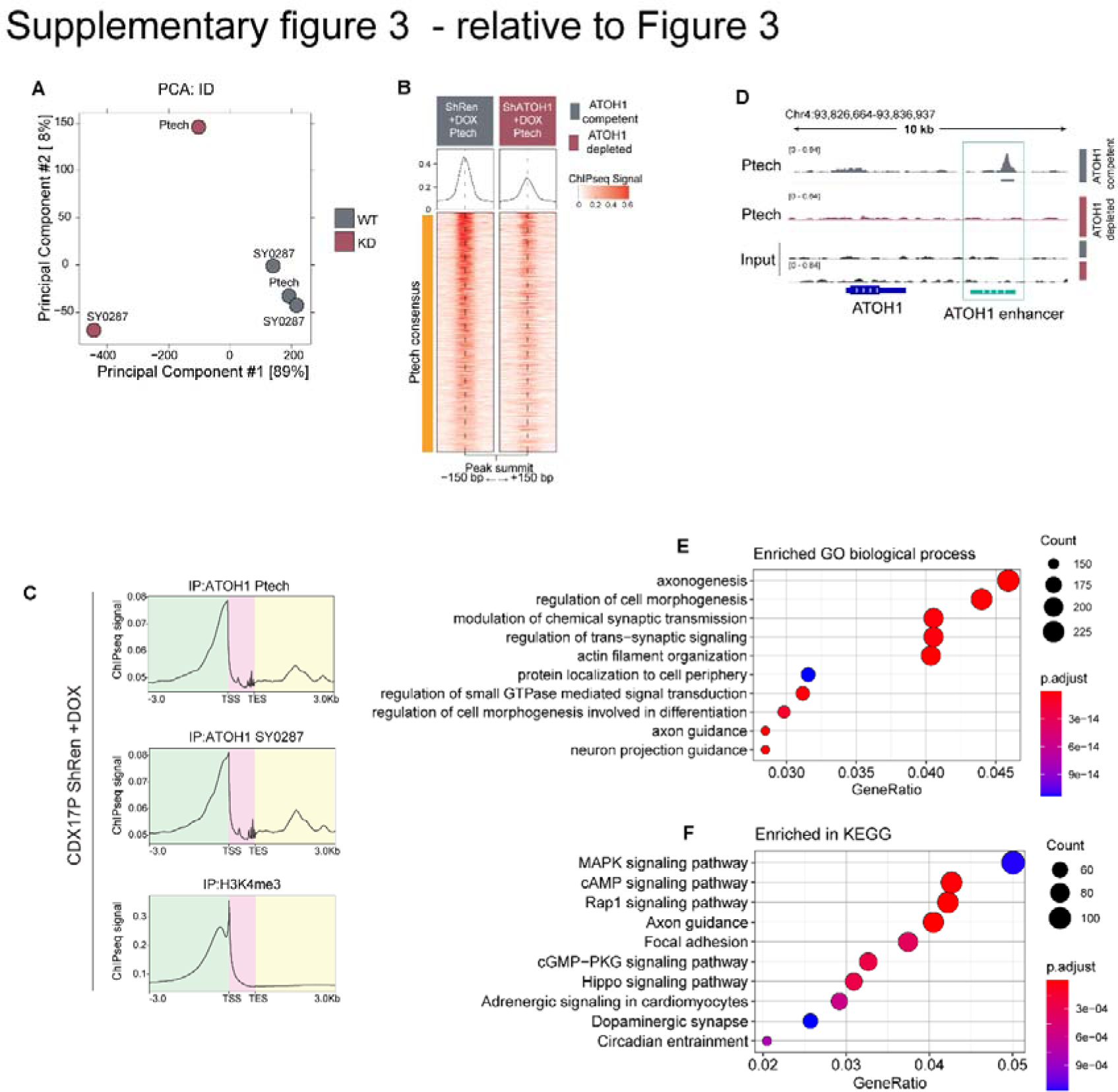
ChIP-Seq samples cluster based on ATOH1 competency and ATOH1 binds to its own enhancer. Relative to. Figure 2. (A) Principal component analysis (PCA) of ChIP-Seq samples where ATOH1 competent samples (grey, WT) cluster together and away from ATOH1-depleted samples (red, KD). (B) Heatmap of ChIP- Seq signal for consensus peak sets of Ptech in ATOH1 competent (grey) and depleted (red) CDX17P, generated with the generateEnrichedHeatmap function within profileplyr v1.8.1^100^. (C) Metagene analysis of ATOH1 (detected with Ptech and SY0287) and H3K4me3 ChIP-Seq signal generated with deepTools^104^. Key: green, upstream of gene body; pink, gene body; yellow, downstream of gene body. (D) ATOH1 binding peaks at ATOH1 locus as detected by the Ptech antibody at the ATOH1 downstream enhancer (light green), which are lost upon ATOH1 depletion. The peaks were visualized with the Integrated Genomics Viewer genome browser. (E-F) Gene ontology (GO) biological process (E) and KEGG (F) enrichment analysis of differentially bound ATOH1 peaks identified Figure 3C-i. Analysis was performed with gage^105^.

**Figure S4.**
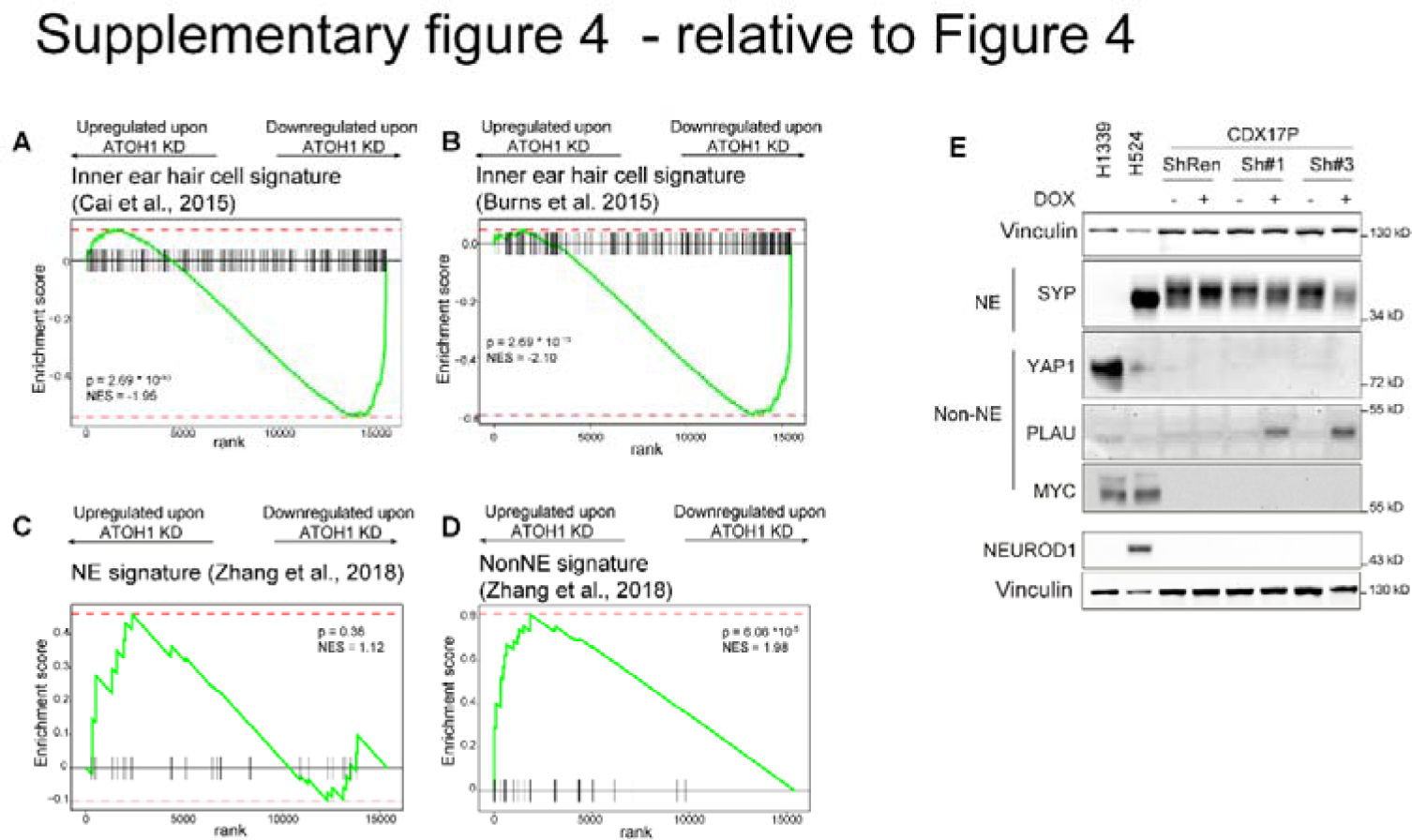
ATOH1 direct targets identified in CDX17P are upregulated in ATOH1 CDX. Relative to. Figure 4. (A-B) Gene set enrichment analysis (GSEA) for inner ear hair cell gene signatures obtained from ref^51^ (A) and ref^52^ (B) upon ATOH1 depletion in CDX17P, performed with Fgsea^103^. (C-D) GSEA for NE (C) and Non-NE (D) gene signatures obtained from ref^106^. NES: normalized enrichment score. (E) Western blot expression of NE marker SYP and NonNE markers YAP1, MYC and PLAU after 14 days of ATOH1 knockdown (KD) induction with doxycycline (DOX) in CDX17P. ShRen served as control for ATOH1 KD; H1339 and H524 served as positive controls for expression of YAP1 and MYC; Vinculin served as loading control. Western blots are representative of N=2 independent experiments.

**Figure S5.**
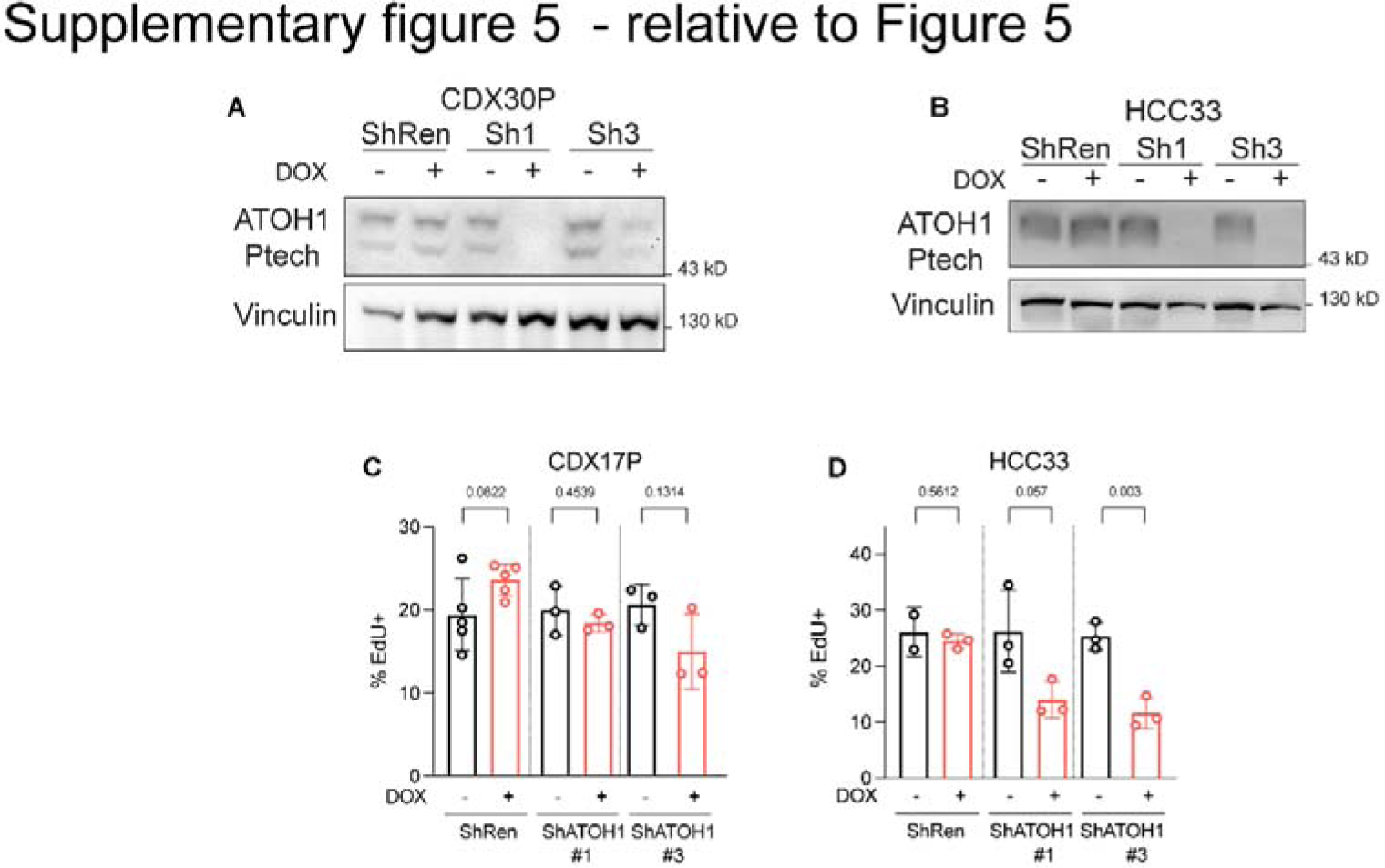
ATOH1 knockdown in CDX17P, CDX30P and HCC33. Relative to. Figure 4. (A-B) Representative western blot for ATOH1 in CDX30P (A) and HCC33 (B) cells transduced with ShRenilla (ShRen) and ShATOH1#1 and #3 and treated with DOX for 7 days. (C-D) Bar plot of percentage of cells in S phase, as identified by EdU incorporation, in CDX17P (C) and HCC33 (D) upon ATOH1 depletion. Statistics are reported as two-tailed unpaired *t* test between DOX untreated and treated condition.

**Figure S6.**
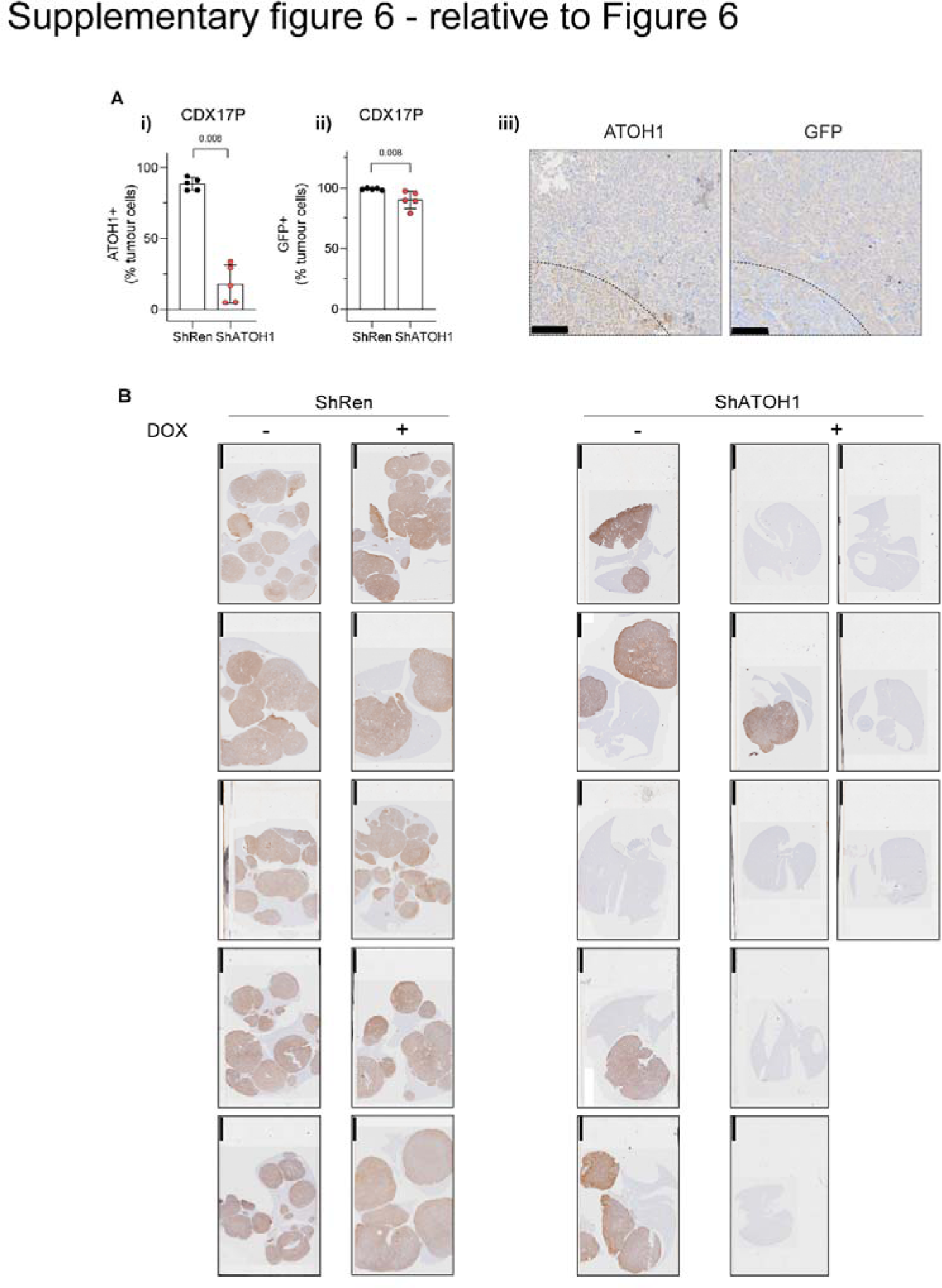
Heterogeneous GFP and ATOH1 expression in ATOH1 KD subcutaneous tumours. ATOH1 KD cells exhibit reduced metastatic ability. Relative to. Figure 6. (A) Quantification of ATOH1 (A-i) and GFP (A-ii) IHC staining in N=5 subcutaneous tumours from mice implanted with either ShRen or ShATOH1 cells and fed DOX-supplemented diet. KD cohort highlighted in red. Statistics reported as per two-tailed unpaired Mann Whitney U test. (A-iii) Representative images of ATOH1 and GFP IHC staining in consecutive sections highlighting parts of tumours negative for GFP and positive for ATOH1 (dotted lines). Scale bars: 100 μm. (B) IHC staining of human mitochondria in livers from animals that underwent intracardiac implantation of ShRen cells and fed a standard diet (-DOX, N=5) or a DOX-supplemented diet (+ DOX, N=5) or ShATOH1 cells and fed a standard diet (- DOX, N=5) or a DOX-supplemented diet (+DOX, N=8). Only one animal in the ATOH1 KD cohort developed metastasis in the liver. Scale bars: 5 μm.

